# Children with Autism Show Impaired Oculomotor Entrainment to Predictable Stimuli

**DOI:** 10.1101/2025.09.03.673494

**Authors:** Shlomit Beker, Oren Kadosh, John J. Foxe, Sophie Molholm, Yoram S. Bonneh

**Affiliations:** Department of Psychiatry, Icahn School of Medicine at Mount Sinai, NY, NY; Department of Pediatrics, Albert Einstein College of Medicine, Bronx, NY; School of Optometry and Vision Science, Faculty of Life Sciences, Bar Ilan University, Ramat Gan, Israel; The Ernest J. Del Monte Institute for Neuroscience, Department of Neuroscience, University of Rochester School of Medicine and Dentistry, Rochester, New York; The Leslie and Susan Gonda Multidisciplinary Brain Research Center, Bar-Ilan University, Ramat Gan, Israel

## Abstract

Individuals with Autism Spectrum Disorder (ASD) show altered synchronization with external events, which may underlie the rigidity and reduced adaptability that characterize the condition. We previously demonstrated that electroencephalography (EEG) recorded from children with ASD reveals impaired neuronal entrainment to predictable visual sequences. Whether similar effects are reflected in other physiological signals remains unclear. Here, we investigated whether eye movement and pupil dilation responses exhibit comparable entrainment differences in ASD. Microsaccades (MS) and pupil diameter were recorded from 31 children with ASD (6-9 years) and 21 age- and IQ-matched typically developing (TD) controls during a task in which four equally spaced visual cues preceded an auditory target. We analyzed modulation of MS release time (MS RT) and pupil response time (pupil RT), along with trial-by-trial variability, as indices of ocular entrainment. Both groups exhibited periodic oculomotor responses to the cues, including phasic MS inhibition and repeated pupil constrictions. In TD children, MS RT and pupil RT increased across cues while their variability decreased, consistent with progressive temporal alignment. These effects were significantly reduced in the ASD group. Oculomotor entrainment measures correlated with EEG inter-trial phase coherence (ITPC) in TD but not ASD children. They also correlated with behavioral response times in both groups, and moderately correlated with autism severity scores. Children with ASD thus showed diminished oculomotor modulation and greater variability in response to predictable stimuli, paralleling earlier EEG findings. These results suggest convergence across physiological systems in indexing impaired processing of predictability in ASD and highlight the promise of multimodal approaches.

## Introduction

Autism Spectrum Disorder (ASD) is characterized by rigid routines and restricted interests, and atypical social communication and interaction. Reduced synchronization of neural oscillations is widely reported in autism^1–4^. A recently forwarded hypothesis links altered synchronization in ASD to abnormal perception and performance^5–7^, and to altered social interactions ^8,9^. This is of relevance to the perspective of ASD as a disorder of adaptation and prediction^10–12^, due to the functional importance of rhythmic oscillations in orchestrating adaptive behaviors, attention and precise action ^13^. We have previously shown evidence for reduced oscillatory entrainment, measured by EEG, in children with autism presented with 4 serially presented visual stimuli, followed by an auditory target ^4^. Although the immediate sensory-evoked responses to the visual stimuli were comparable between the groups, the ASD group showed impaired cortical anticipatory activity prior to the cues, and reduced neural entrainment, measured by inter-trial phase coherence (ITPC), suggesting reduced synchronization between phases of the brain oscillations, and external temporal events.

Entrainment is a widely observed phenomenon by which diverse biological systems adjust their dynamic fluctuations through interaction with the external stimuli. Neural entrainment couples rhythmic brain activity to rhythmic signals, and thus dynamically structures intrinsic neural oscillatory activity to enhance prediction, perception and performance^14–16^. Similarly, eye movement ^17^ and pupil dilation^18,19^ activity couple with repeated sensory stimuli, reflecting cognitive control, attention and anticipation regimes. In the current research we tested whether oculomotor activity of children with ASD, measured by eye movements and pupil dynamics, present a similar pattern of activity to what was found for cortical activity: reduced modulation and entrainment to repeated visual stimuli.

When presented with abrupt visual or auditory stimulus, eye movements are inhibited momentarily in a process known as “oculomotor freezing”^20^ or Oculomotor Inhibition (OMI). This process appears to be a general effect, extending beyond saccades to other eye movements such as blinking^21,22^, ocular drift^23^ and smooth pursuit, including catchup saccades during pursuit^24^. Within OMI research, a particular focus has been placed on microsaccades (MS). These are small, rapid eye movements that occur during fixation, typically at a rate of 1-3 per second and with an amplitude less than 1 degree^25^. While recent studies demonstrate that MS can be produced deliberately^26^ and even used for detailed exploration of a scene^27^, they typically occur involuntarily like spontaneous blinking. Interestingly, the rate of MS decreases significantly in anticipation of repeated stimuli^28,29^, reflecting a form of temporal entrainment to predictable events^30^.

OMI dynamics typically exhibit inhibition followed by rebound. After a stimulus, microsaccade likelihood drops around 150ms (inhibition phase), then rises rapidly, peaking near 400ms before returning to baseline^31^. This time course depends on stimulus characteristics, attention, and anticipation^22,25,32–36^. MS rate is modulated by task difficulty, working memory load^37^, familiarity with names or faces^38^, and speech segmentation^39^. OMI function may be to suppress visual input while processing stimuli and to enhance signal synchronization in the visual cortex^40^.

Abnormal gaze patterns in ASD are often interpreted in the framework of reduced social interest^41^ or a tendency to avoid looking at faces, particularly the eyes^42–45^. Studies have also focused on low-level oculomotor function explanations in ASD children, with mixed results. Some suggested that the basic oculomotor properties, such as saccadic adaptation to simple visual targets^46^ or the number and duration of fixations and saccades when watching animated movies^47^ are comparable between ASD and TD, while others support an idiosyncratic pattern of eye movement^48,49^ and pupil regulation^50^ among ASD individuals. Interestingly, one study found that children with ASD show reduced temporal coherence of saccade and blinking^51^, which was associated with degree of the severity of their ASD symptoms. Together, these findings suggest that while basic oculomotor properties may be intact, the modulation of oculomotor activity by external stimuli is less coherent and more idiosyncratic.

In the current study, we test oculomotor modulation by a sequence of visual stimuli in children with ASD, when the stimuli appear isochronously and followed by an auditory target. We hypothesize that similarly to cortical activity which was recorded in tandem and showed reduced entrainment^4^, ocular activity and ocular entrainment to these stimuli, defined as the variability of microsaccade release time (MS RT SD) and pupil dilation response time (pupil RT SD) across trials, is reduced in ASD. Since correlation between EEG and oculomotor activity in ASD participants is sparsely reported, the novelty of this study lies both in evaluating ocular entrainment in children with ASD to isochronous, predictable stimuli, and in measuring the correlation between entrainment of EEG and oculomotor activity.

## Materials and Methods

### Participants

The data used for this study were collected in the context of a study on the efficacy of different behavioral interventions, in which children diagnosed with ASD were tested in multiple sessions. The data presented here were recorded at the pre-intervention visit. Data from 36 children with ASD between the ages 6-9 years old, and 27 IQ- and age-matched typically developing (TD) children were originally included in the study. Following exclusion due to noisy or insufficient data, 31 children diagnosed with ASD (females: 7; mean age: 7.88 ± 2.5; mean IQ: 97.8 ± 15) and 21 IQ- and age-matched typically developing (TD) controls (females: 12; mean age: 7.77 ± 1.45; mean IQ: 93.6 ± 13) were analyzed. Data collection included recording from a 64-channel EEG, oculomotor (see below) and clinical evaluation. For a full description of participants’ characteristics, clinical evaluations, and acquisition of EEG and behavioral data, see^4,52^.

### Stimuli and Task

The paradigm was designed like a computer game, with stimuli that consisted of a cartoon dog’s face as the visual stimulus. The visual cue stimuli were presented centrally on a 25” ViewSonic screen (refresh rate: 60 Hz, pixel resolution: 1280×x1024×32) using Presentation^©^ software (Neurobehavioral Systems, Inc., Berkeley, CA), and subtending ∼4.4° of visual angle. The auditory target stimulus was a 1000Hz tone 80ms in duration that was delivered at an intensity of 75 dB SPL via a single, from a centrally located loudspeaker (See Figure 1B for paradigm schematic. For a full description of the stimuli see Beker et al., 2021b. The current study presents the oculomotor data. During the paradigm, participants had to respond to a cued / non-cued auditory target as quickly as possible, and then they received feedback to indicate if their response was too fast (e.g., an anticipatory response), right on time, or too slow. The experiment included two conditions: For the *Cue* condition, participants were presented with a sequence of 4 isochronously presented visual stimuli for a duration of 80ms each, presented at a Stimulus Onset Asynchrony (SOA) of 650ms, followed by the auditory stimulus, presented 650ms after the last visual cue. This setup of 4 isochronous visual cues followed by an auditory target was designed to create a highly temporally predictable situation, which in turn was expected to lead to the engagement of neural preparatory processes (i.e., the CNV), and to entrainment to the stimulation rhythm. For the *No-Cue* control condition, the auditory target was not preceded by a sequence of visual cue stimuli. Instead, a central focus “+” sign appeared at the beginning of the trial, and the interval between trial onset (either the central fixation cross or the first of the 4 visual cues) was maintained such that the auditory target appeared 2600ms after the beginning of trial. Both conditions included 15% catch trials on which the auditory target was not presented. Cue and No-Cue conditions were blocked, and presented in runs of 25 trials per block, over a total of 20 blocks. The order of blocks within the experiment for a given participant was chosen randomly prior to each experimental session. Each block lasted 3.5 minutes. Participants were encouraged to take short breaks between the blocks as needed. The whole experiment, including preparation and clean up, lunch, and frequent short breaks, lasted around 3 hours. Participants were seated at a fixed distance of 65 cm from the screen and responded with their preferred hand. In all trials, they were instructed to press a button on a response pad (Logitech Wingman Precision Gamepad) as soon as they heard the auditory tone. Instructions were worded as following: *There will be a cross in the middle of the screen. Try to keep looking at it. Every time you hear a beep, press this button. Press only when you hear the beep and try to avoid pressing when a beep does not occur.* Responses occurring between 150 to 1500ms following the auditory tone were considered correct, and positive feedback was provided via presentation of a cartoon dog image and an uplifting sound. If the response was outside this time window, a running dog cartoon with a sad sound was presented to indicate that the response was too fast, and a sitting dog image with the sad sound was presented to indicate that the response was too slow.

**Figure 1.**
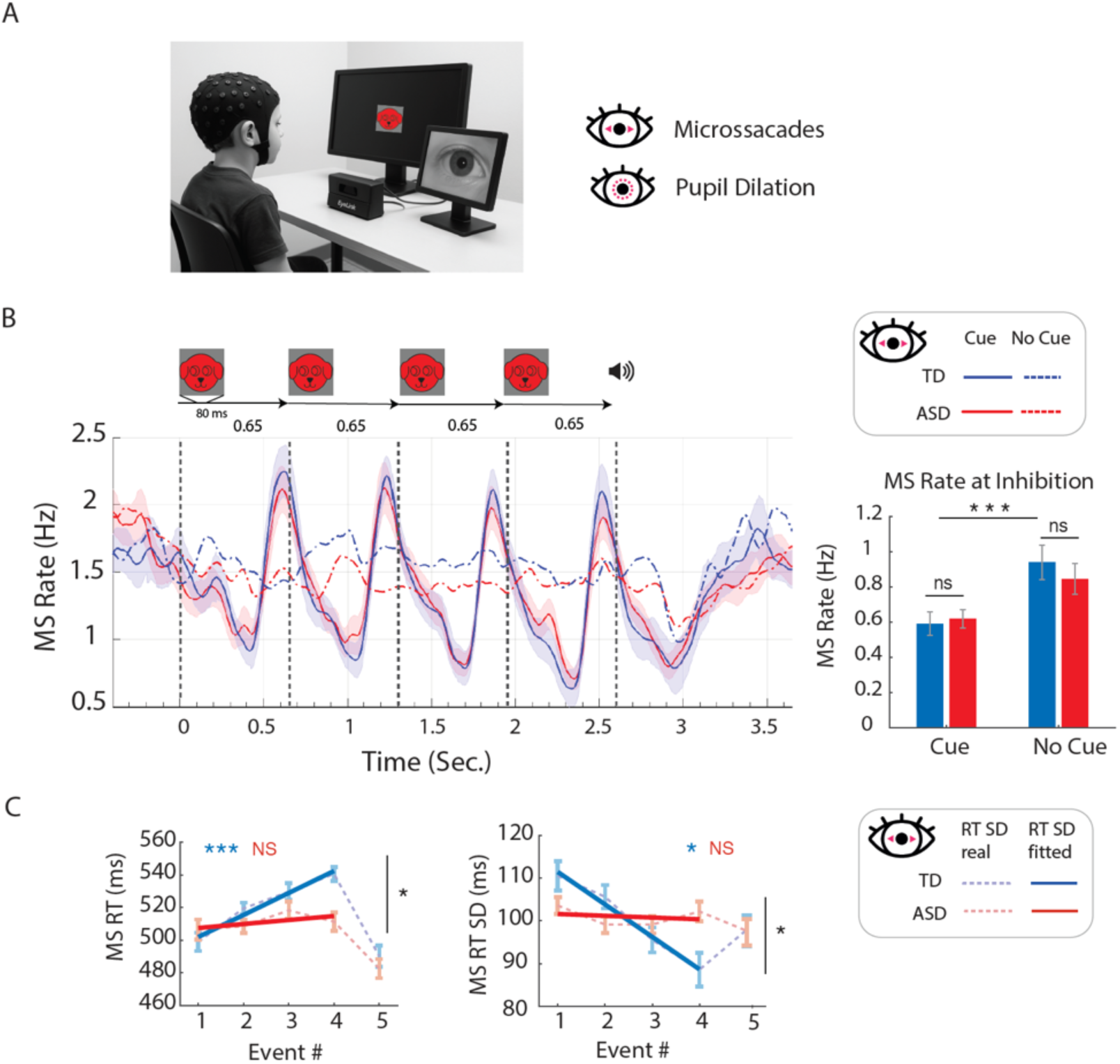
Modulation of microsaccades by the cues. **A.** Left: experimental setup: a child was seated in front of a screen, while EEG (not reported in this paper) and eye tracking data are being recorded simultaneously. (This is an AI-generated photo and does not represent a real human). Right: two oculomotor measures were collected: microsaccades and pupil dilation. **B.** Top: stimuli presented in each trial: 4 visual cues every 0.65 sec (dashed lines) were followed by an auditory target. Bottom: The microsaccade (MS) rate modulation, time-locked to the first cue (time 0). Data is presented for Cue and No Cue conditions. As shown, the microsaccades from both groups were modulated (through microsaccade inhibition and rebound) in response to the visual cues, unlike the no cue condition. Vertical lines: stimulus onset. Right: inhibition of MS (troughs in the graph on the left) shows Cue effect and null Group effect in. **C.** Left: Linear fit of the MS release time (RT) response (average latency of 1^st^ microsaccade in the window 300-650 ms post cue stimulus events). Right: Linear fit of the variability (standard deviation; SD) of the MS response. As shown, MS RT gradually increased while SD decreased with the repeated cues in both groups, but significantly only in the TD group. (*p<0.05; **p<0.01; ***p<0.005).

### Data acquisition

data were recorded with the Presentation^©^ software. EEG data was collected along with Eye tracking (see below) and reaction times. EEG data was analyzed and reported previously ^4,52^. Analog triggers indicating the latencies of stimulus onsets and button presses were sent to the acquisition PC via Presentation® and stored digitally at a sampling rate of 512 Hz, in a separate channel of the EEG and oculomotor data files.

### Eye tracking

eye tracking and pupil dilation were recorded throughout the experiment using the Eyelink1000© eye-tracking system (sampling rate: 1000Hz), and a video camera. Participants were instructed to focus on the center of the screen. However, if the experimenter either noticed through video monitoring that a participant was looking away, or the eye-link system indicated that gaze moved away from the screen, the experimenter reminded the participant to look at the centrally placed fixation cross. We compared the ratio between gaze outside/inside window of 8 visual angels centered at the fixation point.

### Data processing

Oculomotor data were processed and analyzed using custom MATLAB scripts (MATLAB r2022a, MathWorks, Natick, MA).

### Microsaccade detection

For the microsaccade detection, we used eye movement velocity algorithm^32^, that has been used in our previous studies^37,53–56^. The raw data were first smoothed using the locally-weighted plot smoothing (LOWESS) method with a window of *15 ms* to optimize microsaccade detection. This was found to be useful especially with noisy recordings^32^. Eye movements with a velocity range of 8–150°/se*cond,* amplitude range of 0.08-1°, and a duration longer than 9ms were allowed. Eye blinks were detected as in our previous study^21^. We first defined periods with zero pupil size and then extended them by estimating the eyes’ closing and opening time periods, based on the vertical eye movement that typically precedes the blink^56^. Eye tracking data were divided into epochs triggered by the stimulus onset in a range of *-0.5 sec to 5 sec* relative to this trigger, with one epoch per experimental trial. Periods of missing data within an epoch (e.g during an eye blink), were discarded from the analysis with an additional margin of 50ms, without discarding the entire epoch. The rejection rate varied across recordings with a mean of *39% ±18 SD* with no significant difference between groups (ASD: 42% ±17, Typical: 36% ±21, p=0.24, F(1,49)=1.38).

### Microsaccade rate

The MS rate modulation function was calculated as in our previous studies^56–58^, mainly to illustrate the time course of microsaccade occurrences without conducting a statistical analysis. The rate was calculated for the raw onsets which are discrete events. Thus, to visualize their temporal dynamics, convolution with a kernel is a necessary signal processing technique, and it is briefly described here. The rate function was computed for each epoch, by convolving a *500/sec* rate (the eye-tracker sampling rate, ^57^ with a Gaussian window with a sigma value of *50ms* at the time of each microsaccade onset (assuming an estimate of only one microsaccade per sample duration for a sampling rate of *500 Hz*). Finally, the rate functions were averaged across participants.

### Microsaccade and pupil response

For each epoch, the first microsaccade release (MS RT), was computed in different time windows relative to each stimulus onset, including four visual cues and an auditory target stimulus. 300-650ms time window was used following each stimulus onset, to capture only MS that occurred following the cue and avoid corrective saccades that occur immediately after. The maximum pupil dilation was extracted within a time range of *100-650ms* post stimulus onset, and pupil response time (pupil RT) was computed as the latency of that peak. This window was shorter due to the faster pupil dynamics. For each MS RT 1-5 or Pupil RT 1-5, we also calculated the standard deviation within participants, specifically for the Cue condition, for further statistical analysis.

### Statistical assessment

To measure oculomotor entrainment to the rapid visual events, we evaluated the MS RT and Pupil RT, as well as their standard deviations (SD), expecting a gradual decrease in the SD across the four visual cues that precedes the auditory target. Additionally, we plotted the linear fit of these measures across events and hypothesized that the ASD group will exhibit a less negative slope compared to Typical participants. For the statistical assessment of the slope differences across Pupil/MS RT and Pupil/MS RT SD, we employed the Linear Mixed-Effects (LME) analysis^59^. Only the Cue condition was included in the analysis. The MS, Pupil RT, and their SD were used as the dependent variable in separate models. Both models included multiple predictor variables: ‘Group’, ‘Event’, and their interaction, and additionally random intercepts of participants and random Event # within participant:

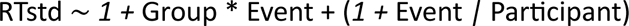

We extracted the slopes and the p-value of the Wald test^60^ for the LME coefficients estimating the Group*Events interaction differences. To ensure more accurate p-values for the fixed effects, Satterthwaite’s method^61^ was used to approximate the denominator degrees of freedom for F-tests and t-tests.

Correlations: we used Pearson to calculate correlations between measures in the study, as followed:

<u>I. Cor</u>ti<u>cal ITPC-oculomotor correlation</u> – We calculated the correlation between EEG measure (ITPC, see^4^ and oculomotor activity (MS and pupil dilation). Of note, one extreme values outliers of the MS RT SD slope excluded from the TD group, msRT SD slope of below -40. Two participants were missing from the EEG recordings, one from each group. (Note that not all EEG participants had oculomotor – 3 didn’t).
<u>II. Motor – oculomotor correlation</u>: we calculated the correlation between motor behavior (button presses at auditory target detection, see^4^ and oculomotor activity, including MS RT, pupil RT and pupil dilation.
<u>III. Clinical scores – oculomotor correlation</u>: we calculated the correlation between oculomotor measures (microsaccades: MS RT slope, MS RT for the 4^th^ Cue, MS RT SD slope; pupil response: Pupil RT slope, Pupil RT for the 4^th^ Cue, Pupil RT SD slope) and the autism diagnostic test scores (autism diagnostic observation scale, ADOS^62^. Since ADOS scores are of a small scale (1-7) and thus highly sensitive to influence of outliers, we excluded data points that exceed ±2 SD from the group mean.
<u>IV. Multiple correlations</u> (for both groups pooled together): we calculated correlation between all measures used in this study. Due to the inclusion of multiple correlations, we applied correction to false discovery rate (FDR)^63^.

## Results

We investigated the effect of isochronous and predictable repeating stimulus on the oculomotor activity in children with ASD. First, we excluded the possibility of a baseline difference in gaze patterns between ASD and TD: We found no evidence for between-groups differences in gaze pattern (mean±std: TD=1.4%+1.2%; ASD=2.1%+1.5%; t=1.6; df=49; p=0.11).

Rate of MS is modulated by the 4 cues similarly for ASD and TD with no Group difference in average rate for MS inhibition for either Cue (F = 0.05(1,54); p = 0.8) (Figure 1B left). Cue effect was significant, with both groups show higher inhibition (lower MS rate) in No Cue vs. Cue condition (F=89(88180,1); p<0.001), and Cue*Group is significant (F=5(88102,1); p=0.02) (Figure 1B, right).

### Microsaccade release time (MS RT) and standard deviation (MS RT SD)

There was no significant group difference in MS RT (LME: F=1.08; df(1,45.8); p = 0.3). However, a strong group x cue interaction was found (LME: F=5.31; df(1,43.2); p = 0.02), implying a weaker modulation of MS RT by cue order in the ASD group. MS RT following each visual cue shows a stronger modulation by the order of the cue in the TD group (slope = 6.5; p = 0.001), but no modulation in the ASD group (slope = 1.5; p = 0.16), and a significant group difference in modulation (LME: F=5.3; df(1,43.2); p = 0.026) (Figure 1C, left).

We then calculating the variability of MS RT, measured by standard deviation of the microsaccade reaction time (MS RT SD). When collapsed across cues, there was a significant group difference in MS RT (LME: F=8.3; df(1, 49); p = 0.005). Again, weaker modulation by cue order was seen in the ASD group (slope = -0.7; p = 0.27), compared with the TD group (slope = -4.3; p = 0.01) and a significant group difference in modulation (LME: F=8.9; df(1,49); p = 0.004) (Figure 1C, right).

### Pupil size modulation

Pupil dilation change was higher for the ASD group compared with TD, although this did not reach significance for either Cue or No Cue conditions (F=0.78 df(1,51764); p=0.37). However, a significant Group*Cue interaction (F=5.9; df=1,51764; p=0.015), meaning that pupil change with higher for the ASD more in the No Cue (Figure 2A).

**Figure 2.**
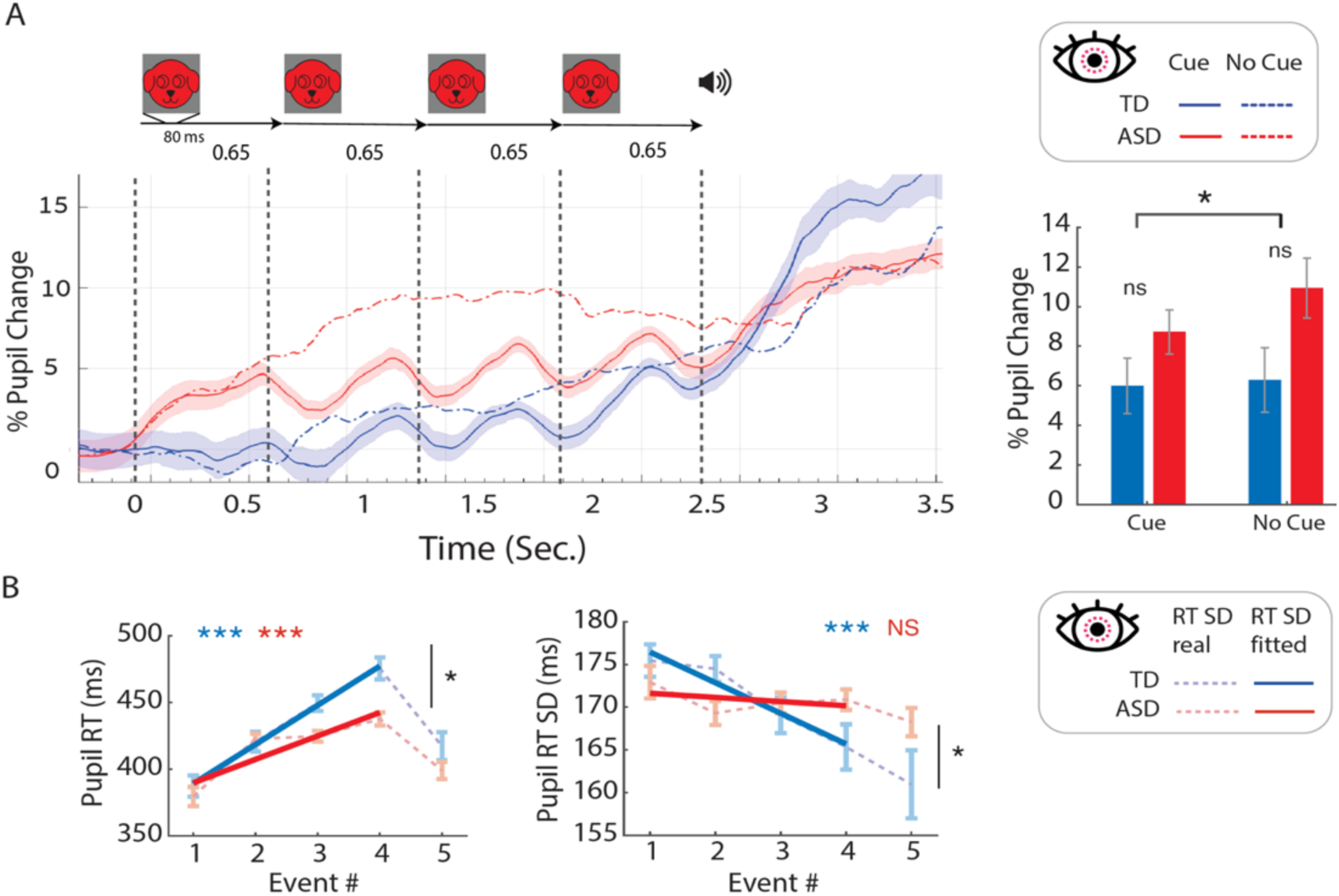
Entrainment of pupil dilation (size) by the cues. **A**. Top: stimuli presented in each trial. Bottom: pupil size modulation, time locked to the first cue (time 0), for the two groups and the two cue conditions (Cue and No Cue), averaged across participants. Pupil size was modulated (light response) for the four cues, but not for the No Cue condition; Right: % pupil change across cues shows no effects for Group or Condition, with a significant Group*Condition interaction. **B**. Left: Pupil RT, calculated as the average latency of the peak pupil size in the window 100-650 ms post stimulus events of each of the 4 cues, had significant different modulation by cue order between the groups. Right: The variability (SD) of the pupil RT, and a linear fit across the 4 cues. Once again, a different modulation is shown for the groups: the variability decreased with the repeated cues for the TD group but did not change in the ASD group. (*p<0.05; **p<0.01; ***p<0.005).

Like MS, modulation of pupil RT, measured as the peak pupil size at 100-650 ms after each cue, and of its variability (pupil RT SD) by the order of the cue was different between the groups (Fig. 2B). Pupil RT for TD: slope=21.1; p=3.89e-14. ASD: slope=13.44; p=4.32e-09. Similarly to MS RT, modulation of pupil RT with cue order was different between the groups (F=4.5; df=1,47.4; p=0.038), but across cues, there was no group difference (F=0.6; df=1,43; p=0.44). Pupil RT SD was significantly higher for ASD than TD (F=4.2; df=1,49; p=0.04), and modulation was significant for the TD (slope=-3.6; p=0.002), but not for ASD (slope=-0.5 p=0.59). A similar pattern is observed across cues: F=5.9; df=1,49.1, p=0.018.

### Cortical - Oculomotor correlation

To test if EEG and oculomotor activity are correlated, we measured linear correlation between the trends (slope) of oculomotor data across the visual cues, represented by modulation of the variability of MS and of pupil dilation latencies. We use these slopes as an indication for entrainment, since the decrease in SD imply better temporal alignment with external stimuli.

Correlation was calculated between cortical oscillatory ITPC (at occipital electrode) and oculomotor activity. The ITPC and the slope of microsaccade reaction time standard deviation (MS RT SD) showed a significant effect in TD, where negative MS RT SD slope (indicating strong modulation on cue order, i.e entrainment) is correlated with higher cortical ITPC values (rho=-0.6, p=0.01). However, this effect was not significant in the ASD group (rho=-0.001, p=0.99). For all participants, there was no correlation (rho=-0.25, p=0.08).

Cortical ITPC and pupil dilation RT SD slope showed inverse trend (All: R=-0.25, p=0.08, N=48), where decreasing SD of pupil response times across 4 visual cues correlated with increasing synchronicity of cortical oscillations. Both groups showed a negative trend (TD: rho=-0.34, p=0.15; ASD: rho=-0.15, p=0.45) (see Figure 3).

**Figure 3.**
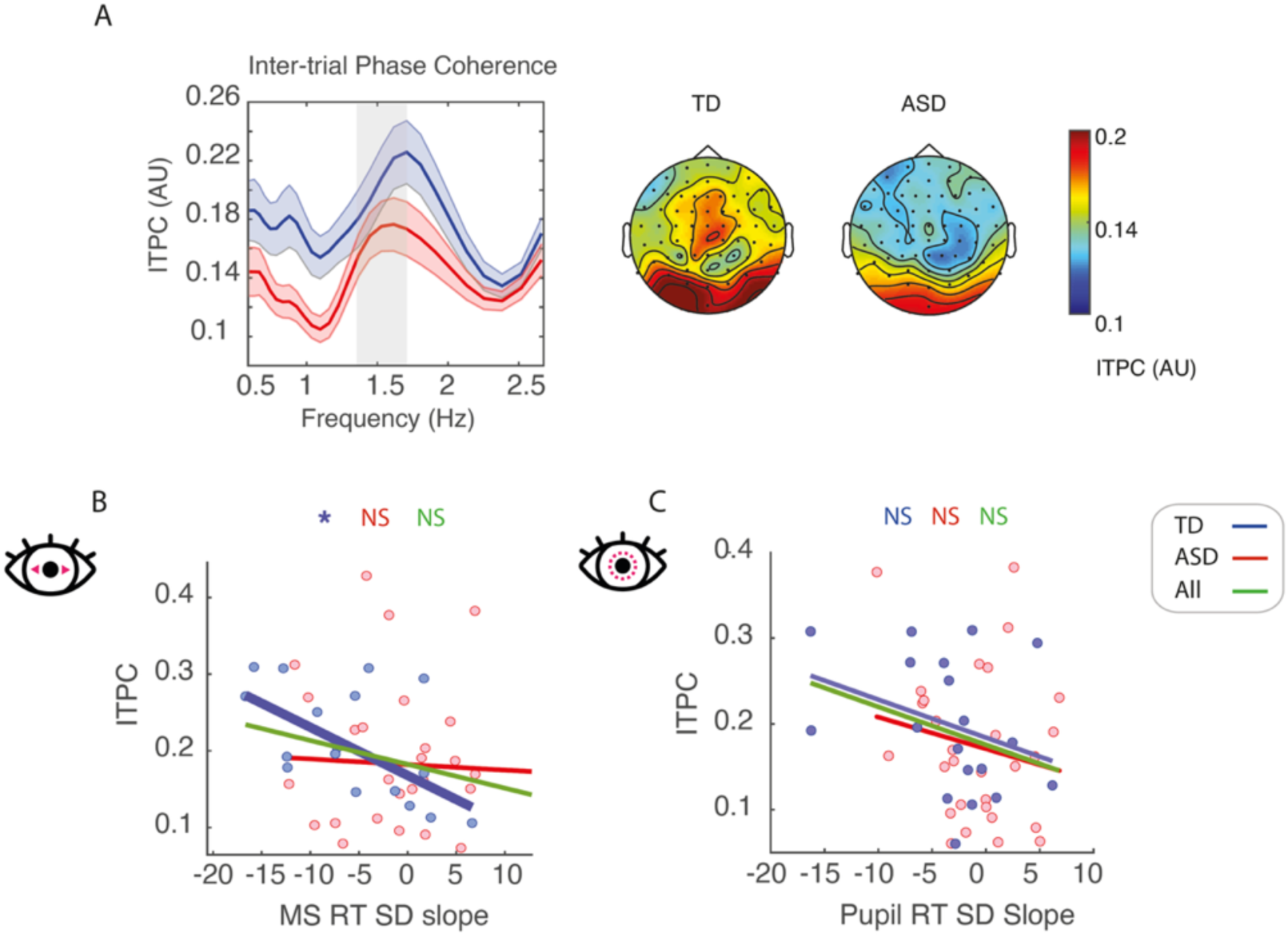
Correlation between oculomotor entrainment and EEG entrainment. **A**. Left: Cortical Inter-trial Phase Coherence (ITPC) as measured by EEG are higher for the TD group around stimulation frequency, 1.5Hz. Right: ITPC topography maps of ITPC at 1.4-1.6Hz show lower ITPC values at occipital and central regions in ASD. **B**. Correlation between microsaccade entrainment, measured as the modulation (slope) of microsaccade reaction time standard deviation (MS RT SD) slope, and cortical ITPC was significant only within the TD group, but not ASD or when both groups collapsed together. **C.** correlation between pupil entrainment, measured as the slope of pupil reaction time standard deviation (Pupil RT SD) slope, and cortical ITPC. This shows a trend for all participants pooled together (p=0.06).

### Motor – Oculomotor correlation

correlations between motor response to the last cue before the target (button presses, as reported before in Beker et al., 2021b), and oculomotor activity show an overall strong coupling between the two types of motor activity in the TD group, and less so in the ASD group (Figure 4), with the exception of Pupil RT, which shows null effect for TD and ASD, but a significant negative correlation for both groups collapsed together. Longer reaction times of MS (MS RT) and change of pupil dilation (Pupil RT) are correlated with shorter motor reaction times, and the degree of pupil dilation following the last cue was positively correlated with reaction times, indicating stronger inhibition for targets with fast motor response. MS RT: *TD: R=-0.42, p=0.074, n=19/20*, *ASD: R=-0.30, p=0.11, n=29/31*, Collapsed: *R=-0.39, p=0.0061, n=48/51*. Pupil dilation peak: *TD: R=0.58, p=0.011, n=19/20*; *ASD: R=0.33, p=0.085, n=29/31*; Collapsed: *R=0.54, p=0.0001, n=48/51*.

**Figure 4.**
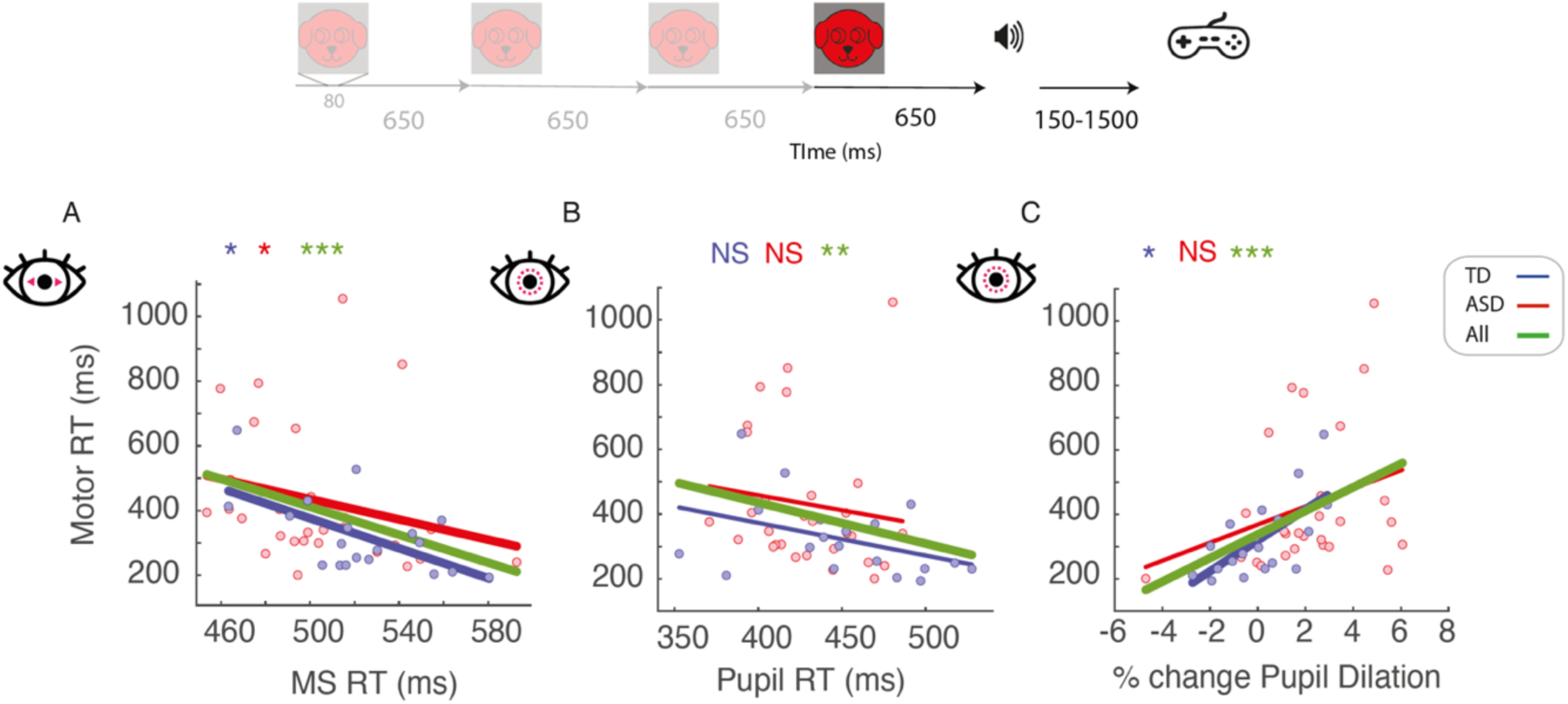
Motor-oculomotor correlations. Top: 4^th^ visual cue precedes the auditory target, to which participants responded. Correlations are calculated between oculomotor responses to the 4^th^ cue, and motor reaction time (rt, in milliseconds, ms) upon target detection. TD (blue), ASD (red) and collapsed across both groups (green), respectively, are presented. **A.** MS RT and motor RT correlation. **B.** Forth cue pupil RT and motor RT correlation. **C.** Pupil dilation peak magnitude and motor RT correlation.

**Figure 5.**
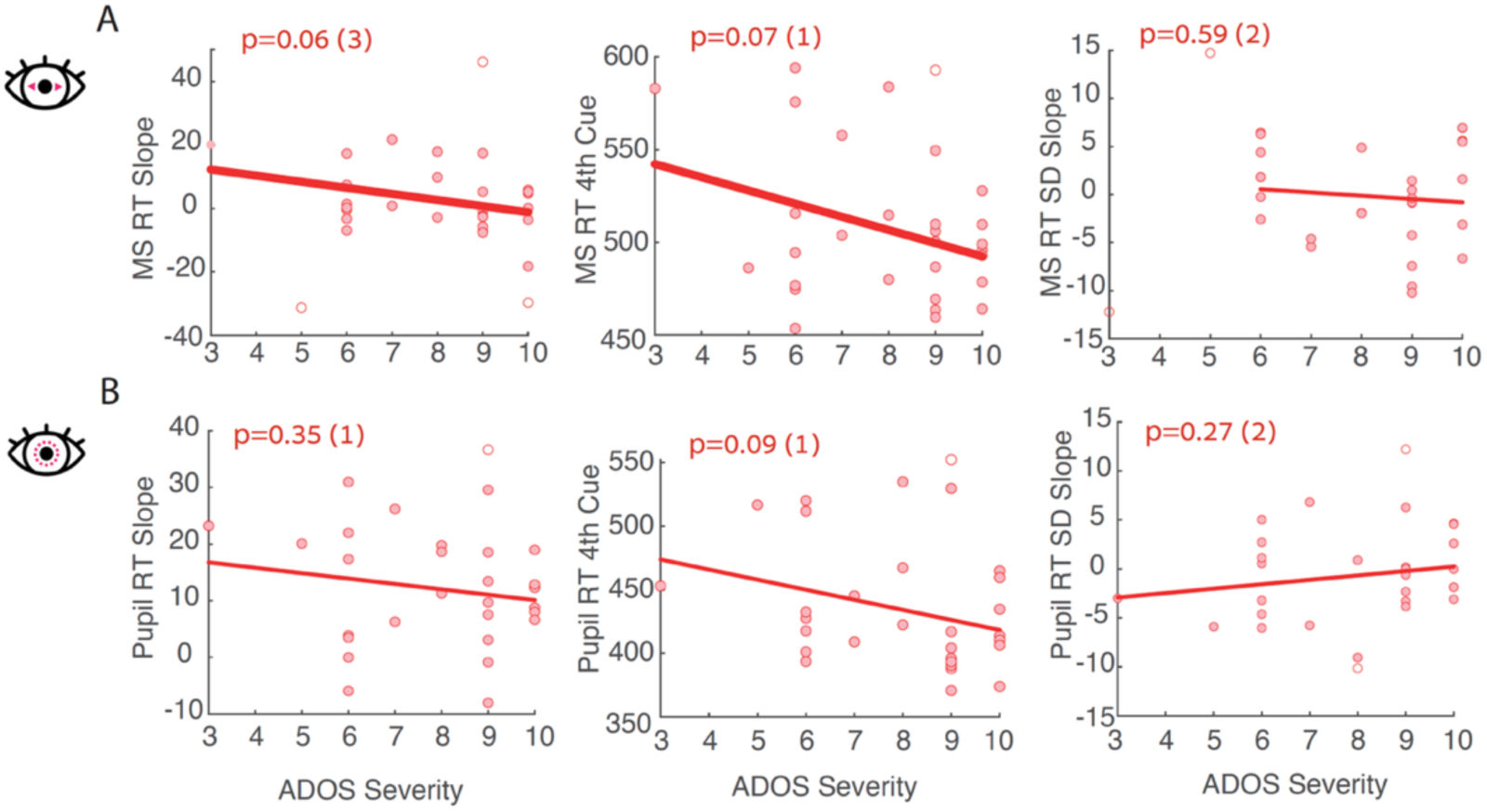
Oculomotor activity - ADOS scores. **A.** Microssacade measures. **B.** Pupil dilation measures. Of all correlations, microsaccade reaction time modulation (MS RT Slope), and microssacade response time to the last cue (MS RT Cue 4^th^) were the only near-significant ones. In parenthesis: number of participants exclude from the correlation analysis, due to exceeding ±2 standard deviations from the group mean. values > 2SD excluded from the correlation analysis.

### Clinical - Oculomotor correlation

correlations between scores from the autism diagnostic observation schedule (ADOS) and oculomotor activity show trend for microsaccade measures: MS RT slope (r=-0.36; p = 0.06) and MS RT 4^th^ cue (r=-0.34; p = 0.07). None of the pupil measures, or the MS RT slope SD is significant. In each analysis, data from between 1 to 3 participants (of N= 30) were excluded due to exceeding ±2 standard deviations from the group mean.

## Discussion

Altered temporal prediction and impaired neural synchronization have been increasingly recognized as important features of autism spectrum disorder (ASD)^5,6,64^. Accumulating evidence suggests that individuals with ASD exhibit reduced neural entrainment ^3,4,65^, contributing to atypical sensory processing, motor planning, and broader difficulties in adapting to structured environmental input ^6,7,10,66^. However, much less is known about whether similar disruptions extend to peripheral physiological activity, such as the oculomotor system (but see ^12^). Our findings demonstrate that children with ASD exhibit reduced oculomotor entrainment to a sequence of visual stimuli in a predictable pattern, consistent with previously reported EEG results in the same participants^4^. This implies that oculomotor activity carries important information, which might overlap with that from EEG.

We tested whether, similarly to effects observed in EEG and in motor performance, entrainment of oculomotor activity is altered in ASD. We focused on the latency dynamics of microsaccades (MS) and pupil dilation, physiological responses that are strongly modulated by attention and arousal, as well as their standard deviation, or inter-trial variability, across repeated trials, as an indication of entrainment. In typically developing (TD) children, as expected ^29,67,68^, we observed greater attention and arousal to be associated with larger pupil dilation, longer response latencies, and reduced variability of these two across stimuli in the trial. However, in the ASD group, this relationship was weaker: hyperarousal was not as correlated with higher latencies, and variability did not change significantly, suggesting a different arousal-attention coupling and reduced oculomotor entrainment overall.

Although both ASD and TD children showed periodic oculomotor responses to the rhythmic visual cues (Figures 1a and 2a), these responses were significantly more variable in the ASD group. Variability of both microsaccade inhibition release latencies (MS RT SD) and pupillary response latencies (pupil RT SD) showed a gradual reduction across successive visual cues in TD children, reflecting increasing temporal alignment with the stimuli (Figures 1b and 2b). This decrease in variability was significantly attenuated in the ASD group, suggesting a diminished ability to adjust oculomotor responses in synchrony with the rhythm of the stimuli. This aligns with the higher variability of oculomotor activity in ASD^51^, as well as with altered predictive temporal processing in ASD^69^. The decreased variability in TD, despite a significant increase in oculomotor latencies, is notable given that standard deviation typically increases with signal magnitude ^70^. The observed delay in oculomotor responses across cue repetitions is also nontrivial, as repeated stimuli are generally associated with motor facilitation through repetition priming ^54,71^. Instead, this delay likely reflects enhanced target anticipation as cue sequences progress, consistent with increased oculomotor inhibition driven by top-down expectations^72^. The significantly reduced preparatory oculomotor suppression observed in the ASD group further supports the notion of diminished predictive processing in autism. One possible explanation is the trend toward greater pupil dilation in the ASD group beginning at trial onset (Figure 2a), which may indicate heightened arousal or hyperfocus^73^. Such an elevated arousal state could interfere with temporal preparation and reduce synchrony with external visual cues.

The correlation between oculomotor measures and EEG interatrial phase coherence (ITPC) in TD further supports the coupling between oculomotor activity and cortical activity: stronger oculomotor entrainment was associated with greater ITPC. This relationship was not significant in the ASD group, indicating a possible decoupling between neural and oculomotor systems. This dissociation may reflect broader deficits in the coordination of sensory and motor processes. The correlation between MS RT and motor behavior (manual reaction times to target) was significant for ASD, suggesting shared temporal control mechanisms. However, motor RT did not show correlation with Pupil RT, whereas it did in TD, supporting the idea that motor planning in ASD may be less tightly coordinated with sensory inputs and anticipatory signals.

Finally, the near-significant correlation between autism severity scores and reduced microsaccade modulation provides some support for the relevance of oculomotor activity to the autism phenotype. Oculomotor data from a larger number of participants is needed to provide stronger support for this association.

Together, our results provide converging evidence that oculomotor modulation by regular sequences of sensory stimuli is disrupted in ASD. This reduced modulation in ASD is consistently associated with impairment across multiple domains, including motor, EEG and the near-significant association with clinical symptom severity, highlighting the relevance of oculomotor activity to the autism phenotype. The correlations between oculomotor modulation with motor response in both groups, and the near-significant association with autism clinical severity underscore the utility of using eye-tracking as a proxy for behavioral response in this population. Further research may explore how these oculomotor markers relate to adaptive functioning and responsiveness to interventions targeting timing and prediction in ASD. Our results extend previous findings of altered anticipatory neural activity in ASD, now showing that oculomotor signals - often considered low-level, reflexive responses - are also less temporally tuned in ASD. Given that MS and pupil responses are sensitive to attention, expectation, and cognitive load, these findings support the notion that temporal entrainment deficits in ASD span across multiple systems.

One seemingly contradictive interpretation can be given to the increasing MS RT and pupil RT with the cue sequence. Highly expected, salient stimuli are found to shorten the inhibition phase^53,74^, while unexpected stimuli prolong it^75,76^. Likewise, even self-induced surprises, like saccade-induced visual transients in oddball paradigms, can lengthen inhibition^37^. According to this, in our settings we could have expected decreasing MS and pupil RT with the cues, which become more predictable. However, while surprise decreases, attention increases, and might have a larger effect on these autonomic dynamics. In this scenario, attentional or arousal resources gradually accumulate in anticipation of the upcoming target.

Finally, not all participants contributed usable EEG and oculomotor data, which slightly reduced statistical power for the EEG-oculomotor correlation. However, the convergence of findings across modalities and analyses lends strong support to the validity of our conclusions.

## Acknowledgment

The authors thank the participants and their families for taking the time to participate in this study and for their dedication to advancing understanding of autism. We acknowledge the contributions of the Human Clinical Phenotyping Core of the Rose F. Kennedy Intellectual and Developmental Disabilities Research Center (RFK IDDRC) and Drs. Juliana Bates and Pamela Counts for performing clinical and cognitive testing.

## Supplementary Material

**Figure S1.**
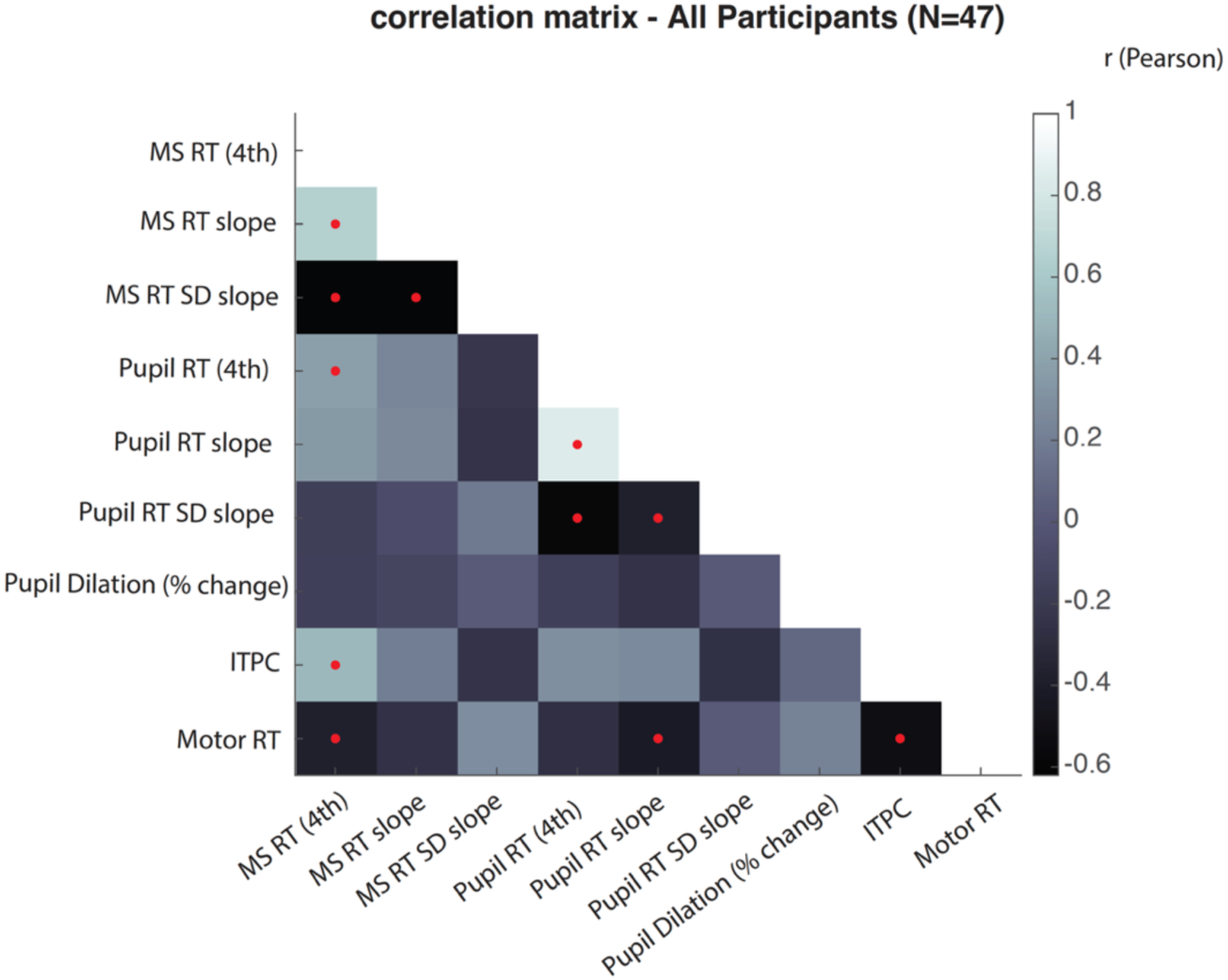
Correlation matrix for all measures, calculated on all participants pooled together (N=47). Grey shades indicate r values (Pearson pairwise correlations). Red dots indicate correlations that survived correction for multiple comparisons.

